# “In Vivo Imaging of Bone Collagen Dynamics in zebrafish”

**DOI:** 10.1101/2023.11.06.563068

**Authors:** Hiromu Hino, Junpei Kuroda, Shigeru Kondo

## Abstract

Type I collagen plays a pivotal role in shaping bone morphology and determining its physical properties by serving as a template for ossification. Nevertheless, the mechanisms underlying bone collagen formation, particularly the principles governing its orientation, remain unknown due to the lack of a method enabling continuous in vivo observation. To address this challenge, we constructed a method to visualize bone collagen by tagging with GFP in zebrafish and observed the interactions between the osteoblasts and collagen fiber during bone formation in vivo. When Col1a2-GFP was expressed under the control of the osteoblast-specific promoters *osx* or *osc* in zebrafish, bone collagen could be observed clearly enough to identify their localization, but collagen from other organs did not. Therefore, we determined that this method was of sufficient quality for detailed in vivo observation of bone collagen. Next, bone collagen in the scales, fin ray, and opercular bones was observed in detail in zebrafish, when bone formation is more active.

By simultaneously observing bone collagen and osteoblasts, we successfully observed dynamic changes in the morphology and position of osteoblasts from the early stages of bone formation. It was also found that the localization pattern and orientation of bone collagen significantly differed depending on the choice of expression promoter. Both promoters (*osx* and *osc*) used in this study are osteoblast-specific, but their Col1a2-GFP localizing regions within bone are exclusive, with *osx* region localizing mainly the outer edge of bone and *osc* region localizing the central area of bone. This suggests the existence of distinct subpopulations of osteoblasts with various gene expression profiles, each of which may play a unique role in the osteogenic process.

These findings would contribute to a better understanding of the mechanisms governing bone collagen formation by osteoblasts.

## Introduction

Type I collagen secreted by osteoblasts aggregates to form a higher-order structure that serves as a scaffold (bone matrix) on which calcium-crystalline hydroxyapatite is deposited. Therefore, many of the physical properties of bone depend on the shape, orientation, and density of the collagen network (Georgiadis *et al*., 2016). Since the collagen orientation can vary from a linear arrangement, in-plane crossing, and layered overlapping, depending on the bone morphology and the direction of force application (Reznikov *et al*., 2014; Georgiadis *et al*., 2016), it is assumed that there is a precise control mechanism. Moreover, since abnormal collagen formation is responsible for pathological conditions such as osteogenesis imperfecta (OI) and various bone formation abnormalities (Marini *et al*., 2007), it is extremely important to elucidate the regulatory mechanisms.

In order to gain insights into how collagen formation and orientation are regulated in bone formation, it is essential to observe the dynamic behavior of collagen and the cells that secrete it in vivo. However, the existing methods used to observe collagen fibers do not fully satisfy this requirement. For example, they rely on fixed samples submitted to X-ray structural analysis, electron microscopy, and reagent staining of bone collagen (Reznikov *et al*., 2014). Some studies have also observed collagen in vivo by collagen-mimetic peptides that bind specifically to collagen (CMP) (Yu *et al*., 2011; Li *et al*., 2012) or by second harmonic generation (SHG) (Campagnola *et al*., 2002; Chen *et al*., 2012; Lebert *et al*., 2015; Lebert *et al*., 2016), but even using these methods, the localization of all collagen in real time isn’t directly reflected, and there are various limitations, such as the size and localization of the target collagen structure and detection sensitivity. Therefore, a new and novel method that allows in vivo observation of the process of collagen formation in bone has been desired.

The most convenient way to visualize molecules in the body is to attach a fluorescent protein such as GFP to the target molecule. Therefore, many groups have labeled Col1 molecules with GFP, but it was difficult to observe clear fiber structures. This is probably because the labeled Col1 molecules inhibit normal aggregation. However, recent efforts to devise binding positions have allowed visualization of bone collagen in cultured osteoblasts and mouse skulls (Lu *et al*., 2018; Shiflett *et al*., 2019). In mice, there are physical limitations to observing whole-body bone collagen, but more comprehensive observations should be possible in more transparent animal species (Morris *et al*., 2018; Kashimoto *et al*., 2022). In this study, we developed transgenic zebrafish lines that expresses Col1a2-GFP in an osteoblast-specific manner and observed bone collagen in vivo during development. Simultaneous observation of collagen and osteoblasts provided insight into how osteoblasts form bone collagen.

## Materials and methods

### 2.1. Zebrafish husbandry

We used AB strain zebrafish for establishing transgenic zebrafish lines. All fish were treated and used in experiments in accordance with the guidelines and approved protocols for animal care and use at Osaka University, Japan. Zebrafish were maintained under standard laboratory conditions at 28.5 °C and a 14/10 h light/dark cycle.

### 2.2. Generation of Zebrafish Transgenic Line

To create *col1a2-gfp* sequence, we modified the zebrafish *col1a2* sequence by replacing the propeptide and telopeptide regions with *gfp* sequences, as previously described by Morris and colleagues (Morris *et al*., 2018). Initially, we generated pGEMT-*col1a2* by cloning the zebrafish *col1a2* sequence from a self-made zebrafish cDNA library into the pGEMT-easy vector using pGEM®-T Easy Vector Systems (Promega). Subsequently, a fragment containing a BamH1 insertion site, was created via mutagenic PCR. The *gfp* sequence was cloned and incorporated using NEBuilder HiFi DNA Assembly Master Mix (NEB). The *col1a2-gfp* sequence from the final pGEMT-*col1a2-gfp* construct was excised using restriction enzymes and inserted into the pTol2 vector, resulting in the creation of pTol2-*osteorix(osx):col1a2-gfp* and pTol2-*osteocalcin(osc):col1a2-gfp* constructs. The *osx* and *osc* promoter regions, known for their osteoblast-specific activity in medaka and also applicable to zebrafish (Inohaya *et al*., 2007; Renn and Winkler, 2009), were obtained from the genome via PCR. For the preparation of the osteoblast visualization transgenic line, we constructed pTol2-*3xosx:lifeact-yfp*, which incorporated the *lifeact-yfp* sequence into the pTol2 vector with three repeats of the *osx* promoter region using NEBuilder. Each plasmid was diluted with double-distilled water (DDW) to a final concentration of 50 ng/ml and injected into 1-cell stage embryos along with 25 ng/ml of Transposase mRNA (Urasaki *et al*., 2006).

### 2.3. Confocal Imaging and image processing

In Fig. 1 and 2A, for whole body imaging, fish were anesthetized using tricaine (MS-222), placed on a glass dish and imaged with a BZ-×710 microscope (Keyence) equipped with a x10 objective (NA 0.45 PlanApo).

**Fig. 1.**
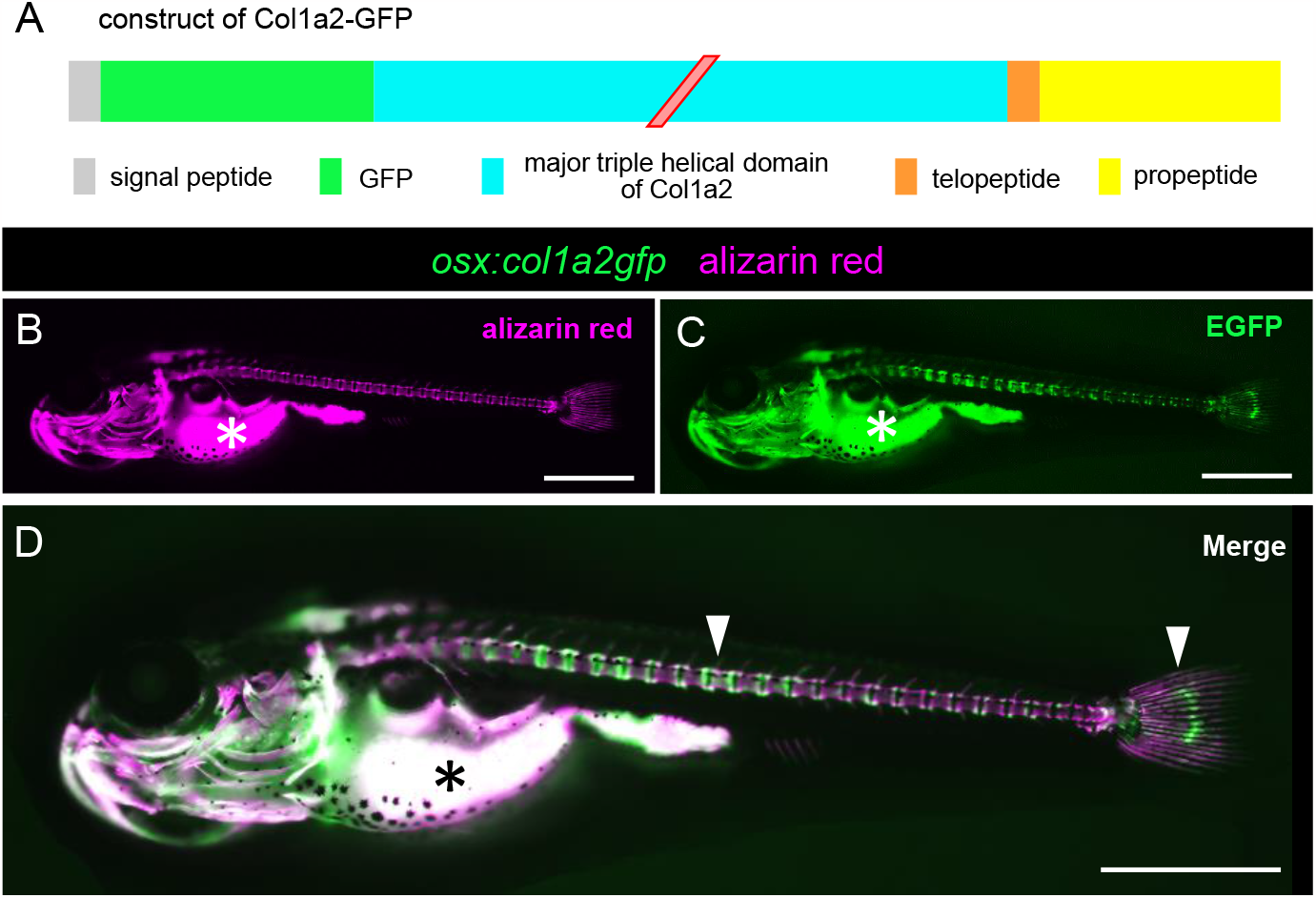
Col1a2-GFP distribution pattern in Tg(*osx:col1a2-gfp*) fish at 14 days post fertilization (dpf) with alizarin red staining. **(A)** A schematic illustration of the expression construct of *col1a2-gfp*. **(B, C, D)** Alizarin red signal (B), GFP signal (C), Merge (D) image of alizarin red stained Tg(*osx:col1a2-gfp*) fish. * in A, B, C shows auto-fluorescence in yolk. Strong accumulations of Col1a2-GFP were observed at specific sites of bones (arrowheads). Scale bar = 1mm.

**Fig. 2.**
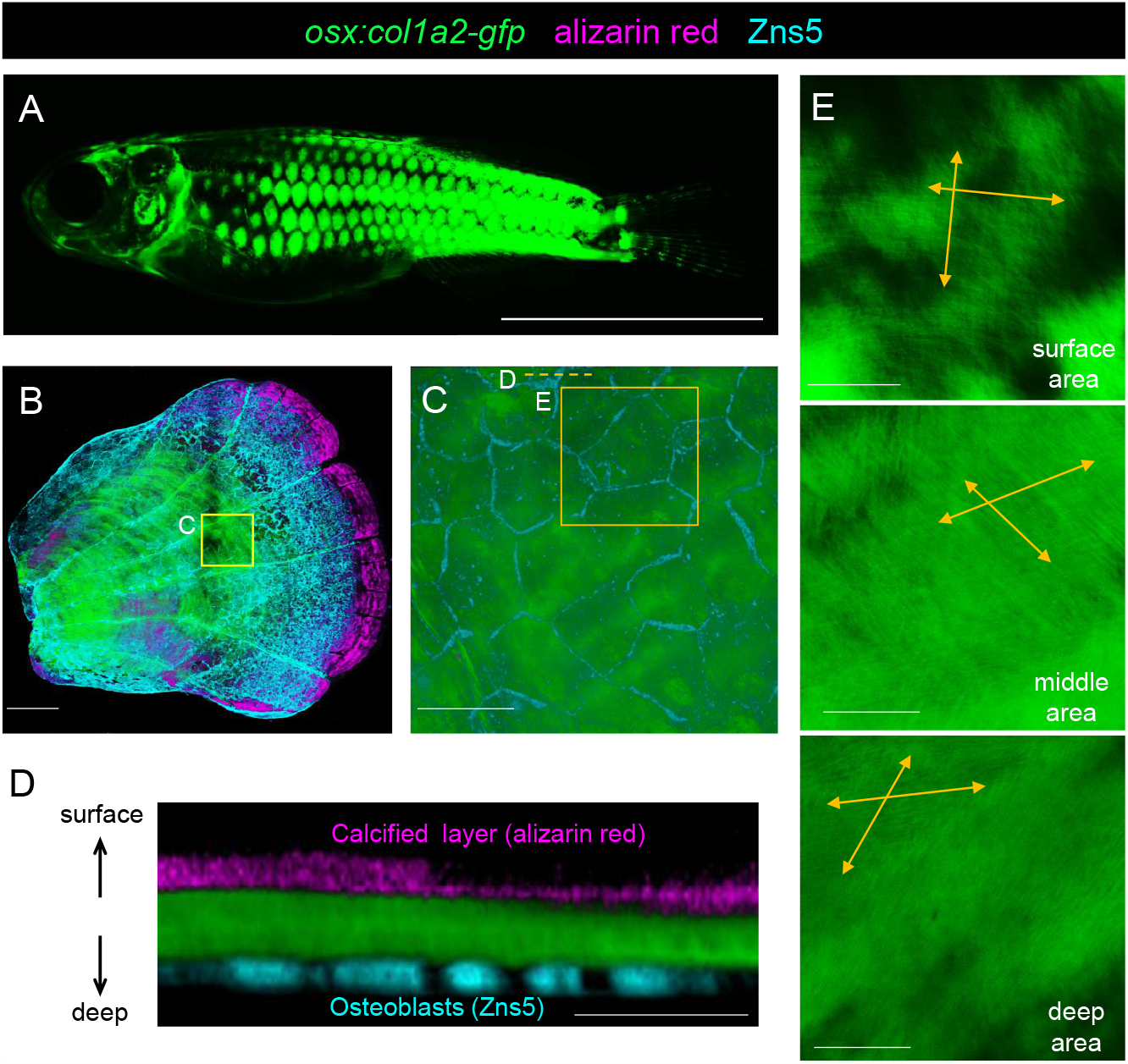
Florescent images of the scales in Tg(*osx:col1a2-gfp*) fish. **(A)** GFP signal of whole body in Tg(*osx:col1a2-gfp*) 2 months old fish. Scales are visualized by GFP florescence. Scale bar = 5mm. **(B)** Whole image of single scale isolated from Tg(*osx:col1a2-gfp*) fish with Alizarin red staining and immunostaining of Zns5 antibody. Scale bar = 200µm. **(C)** Magnified image of yellow box C in (B). Zns5 fluorescence signals show plasma membrane of osteoblasts. Scale bar = 50µm. **(D)** Section view at yellow dotted line D in (C). Col1a2-GFP localizes under osteoblasts. Strong calcification was observed at surface of scale without Col1a2-GFP signals. Scale bar = 60µm. **(E)** Slice images of the scale along their front and back axes in box E in (C). Different oriented collagen fiber indicated by yellow arrow were observed in each image. Scale bar = 20µm.

For fish in which bone images were captured in Fig. 2 B, C, D, E, 3, 4, 5 and Sup. Fig. 1, 2, euthanasia was performed using tricaine, followed by fixation in 4% paraformaldehyde (PFA) in phosphate-buffered saline (PBS) at 4°C overnight. Bone samples were imaged with a confocal microscope, Stellaris8 (Leica) with objectives, HC PL APO 10x/0,40 CS2, HC PL APO 40x/1.30 OIL CS2, HC PL APO 63x/1,40 OIL CS2 (Leica). All images were analyzed using Imaris (Oxford Instruments) and LASX (Leica) software to generate 3D and section images.

### 2.4. Alizarin red Staining

For alizarin red staining, live fish were immersed in a 0.005% alizarin red solution at 28.5°C overnight. Subsequently, the fish were returned to their tank water, allowed to swim for 1 hour for washing, and the samples were collected.

### 2.5. Immunostaining

For immunostaining, samples were washed in phosphate-buffered saline with 0.1% Tween 20 (PBST) after fixation. They were then blocked in PBST containing 1% goat serum at room temperature for 1 hour. Next, the samples were incubated in 1% goat serum PBST with the primary antibody anti-Zns5 (ab181413, Abcam, 1:200) at 4°C overnight. Following a PBST wash, the samples were incubated in 1% goat serum/PBST with Alexa Fluor 633 goat anti-mouse IgG antibody (Invitrogen, 1:200) at 4°C overnight.

## Results

### 3.1. Expression of Col1a2-GFP in Zebrafish Osteoblasts Enables Systemic Bone Collagen Visualization

Lu’s group found a new insertion position that is not inhibitory to collagen molecules polymerization and can be visualized by GFP at telopeptide (Lu *et al*., 2018). We followed their method and bound GFP to the zebrafish *col1a2* gene. We also expressed *col1a2-gfp* under the control of the osteoblast marker gene *osterix*-derived promoter (Renn and Winkler, 2009) to specifically visualize bone collagen (See Fig. 1A for construct details). A whole body image of strain Tg(*osx:col1a2-gfp*) larvae at 14 days post fertilization (dpf) obtained by gene transfer is shown in Fig. 1. The GFP signal was consistent with the alizarin red-labeled bone location, suggesting Col1a2-GFP localization in bone (Fig. 1B, C, D). The background fluorescence was also very low, indicating that collagen other than bone was not labelled by Col1a2-GFP. Because bone collagen-specific labelling was obtained as expected, we next used this strain to examine the later stages of osteogenesis in more detail.

### 3.2. Visualized Bone Collagen Exhibits Normal Fiber Structure

Scales, which correspond to dermal bone, are a valuable resource for bone research due to their location on the surface of the fish body, which makes them easily observable and amputable (Sire *et al*., 1997; Sire and Akimenko, 2004). Given that previous studies have reported the detection of *osx* expression in scales and that electron microscopy observation has revealed a well-ordered, collagen-based layered structure within these scales(Iwasaki *et al*., 2018; Kobayashi-Sun *et al*., 2020), we initiated our investigation by examining the localization of Col1a2-GFP in scales. Fig. 2A is a whole body image of a 2-month-old zebrafish, clearly showing the outline of the scales. When the scales were peeled off and observed directly with a confocal microscope (Fig. 2B), a concentric circular pattern was visible. The fluorescence of GFP was stronger in the center of the scales and weaker at the edges. This could be attributed to the varying levels of *osx* expression in osteoblasts, which seem to depend on their location within the scales.

When the area with strong GFP fluorescence was observed at higher magnification, the collagen fiber structure became visible. The collagen fibers are aligned in a uniform direction, showing characteristic lattice-like pattern reported in previous study (Campagnola *et al*., 2002; Feng *et al*., 2020) (Fig. 2E). We also observed that the orientation of the collagen changed in each layer (Fig. 2D, E) when the focus was changed. These results indicate that secreted Col1a2-GFP is normally incorporated into bone collagen fiber structures and visualizes them.

### 3.3. Localization of Bone Collagen Visualized by Col1a2-GFP Varies with Promoter Dependence

As shown in Fig. 2B, the GFP signal was stronger in the center and weaker at the margins in the scale (Fig. 2B). Furthermore, it also localized under osteoblasts and didn’t in calcified area stained with alizarin red (Fig. 2D). We considered the possibility that this region specificity might depend on the nature of the *osx* promoter and decided to create a Col1a2-GFP construct using a promoter derived from osteocalcin (*osc*), another osteoblast-specific marker gene, to compare their Col1a2-GFP localizing regions. It is known that *osteocalcin* expression is weak in scales, and indeed, fluorescence signal of Col1a2-GFP could not be confirmed in scales of Tg(*osc:col1a2-gfp*) fish. However, Col1a2-GFP in the *osc* promoter strain was observed in the fin ray and opercular bones, both of them are often used for bone research in zebrafish due to ease of observation (Kimmel *et al*., 2010; Marí-Beffa and Murciano, 2010). But Col1a2-GFP distribution in both bones was different from that of the *osx* promoter (Fig. 3 and 4).

**Fig. 3.**
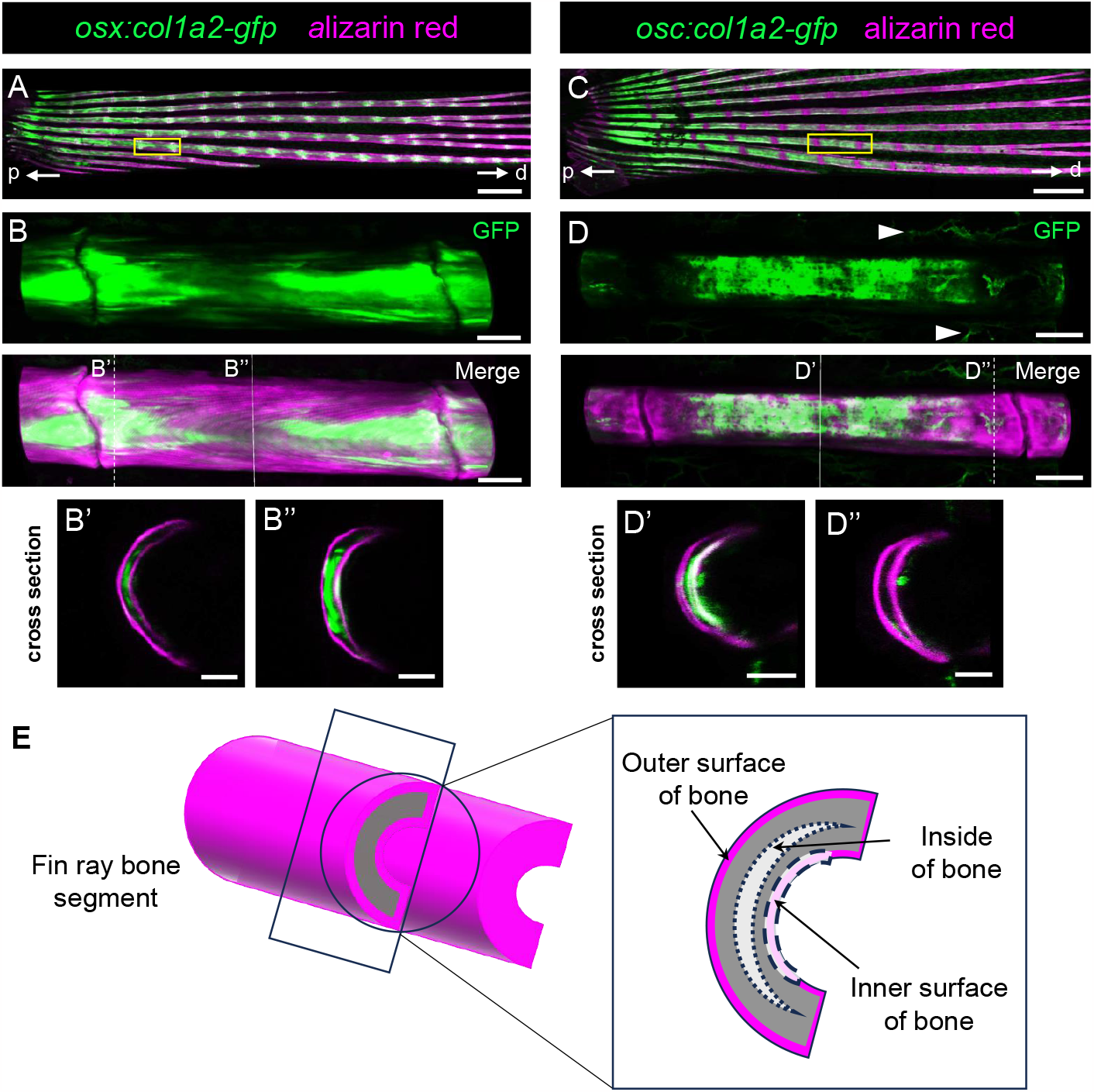
Fluorescent images of the Alizarin red stained fin ray bones in Tg(*osx:col1a2-gfp*) and Tg(*osc:col1a2-gfp*) **(A, C)** Alizarin red stained fin ray bones in Tg(*osx:col1a2-gfp*) (A) and Tg(*osc:col1a2-gfp*) (C). Arrows p, d indicate proximal-distal axis of fin ray bones. **(B, D)** Magnified image of yellow box in (A, C). Arrowheads in (D) indicate auto-fluorescence of blood vessel. (B’, B’’ and D’, D’’) Section views at lines B’, B’’ and lines D’, D’’ in B, D. **(E)** Illustration of fin ray bone segment. Col1a2-GFP driven by *osx* promotor was detected inner side of bone at edge of bone segment (B’’) but detected weakly at middle of bone segment (B’). Driven by *osc* promotor, it was detected at inner surface of bone at middle of bone segment (D’) but detected significant weakly at edge of bone segment (D’’). Scale bar = 400µm (A, C), 40µm (B, D) and 20µm (B’, B’’, D’, D’’).

**Fig. 4.**
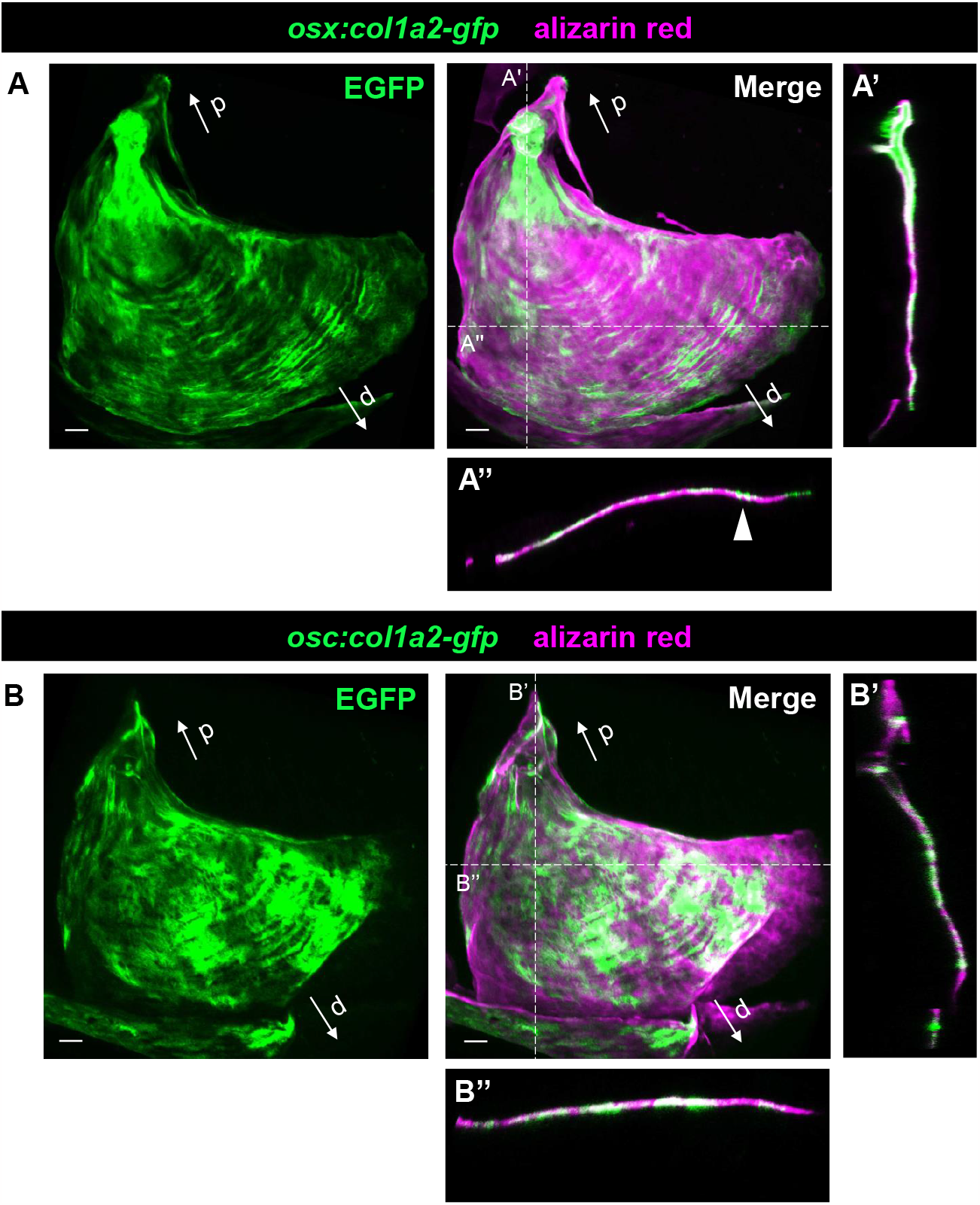
Fluorescent images of the Alizarin red stained opercular bones in*Tg(osx:col1a2-gfp)* and *Tg(osc:col1a2-gfp)* **(A, B)** The opercular bones stained with Alizarin red isolated from *Tg(osx:col1a2-gfp)* (A) and *Tg(osc:col1a2-gfp)* (B) fish. Arrows p, d indicate proximal-distal axis of opercular bones. (A’, A’’) Cross-sectional images at white dotted lines A’, A’’ in (A). (B’, B’’) Cross-sectional images at white dotted lines B’, B’’ in (B). Arrowhead in (A’’) indicates strong accumulation of Col1a2-GFP at limited sites. Scale bar = 20µm.

As shown in Fig. 3A and B, under the *osx* promoter, Col1a2-GFP localized near the articular regions of the fin ray bone. In contrast, under the *osc* promoter, it was observed predominantly at the central part of the fin ray bone (Fig. 3C, D). With the *osx* promoter, collagen fibers tended to align along the long-axis direction of the fin ray bone (Fig. 3B), while no distinct orientation was observed with the *osc* promoter (Fig. 3D). Moreover, the localization pattern in cross-section also exhibited clear differences: under the *osx* promoter, GFP localization was confined to the inside of the fin ray bone (as alizarine red stains only the surface of the fin ray bone), whereas under the *osc* promoter, Col1a2-GFP was primarily found at the inner surface of the curved fin ray bone (Fig. 3B’, B’’, D’, D’’).

In the opercular bone, Col1a2-GFP localization expressed under the two promoters also showed partially overlapping but roughly exclusive patterns. Under the *osx* promoter, strong GFP signals localized in the proximal region and near the outer edge of the fan-shaped opercular bone (Fig. 4A). In contrast, under the *osc* promoter, Col1a2-GFP localized around the central region of the fan (Fig. 4B). In addition, the overall GFP pattern tended to be arranged in an arc-like pattern (Fig. 4A, A’’ arrowhead). This arc-like localization is consistent with the progressive zones of annual growth observed in growth analysis of opercular bone using alizarin red (Kimmel *et al*., 2010). Similar arcuate striations were observed in opercular bone of the Tg(*osc:col1a2-gfp*) fish, but the Col1a2-GFP localizing area differed from that of the Tg(*osx:col1a2-gfp*) fish because they were more extensive (Fig. 4B, B’’).

### 3.4. Simultaneous Observation of Bone Collagen and Osteoblasts Facilitates Osteoblast Dynamics Analysis during Bone Collagen Formation

The opercular bone is one of the first bones to form during development and located near the body surface, making it a suitable model for observing bone formation. To gain insight into osteoblast dynamics and their role in bone collagen formation, we crossed the osteoblast visualization line Tg(*3xosx:lifeact-yfp*) with Tg(*osx:col1a2-gfp*) and observed growth process of opercular bone during early stages of development (3, 5 and 7 dpf) (Fig. 5).

**Fig. 5.**
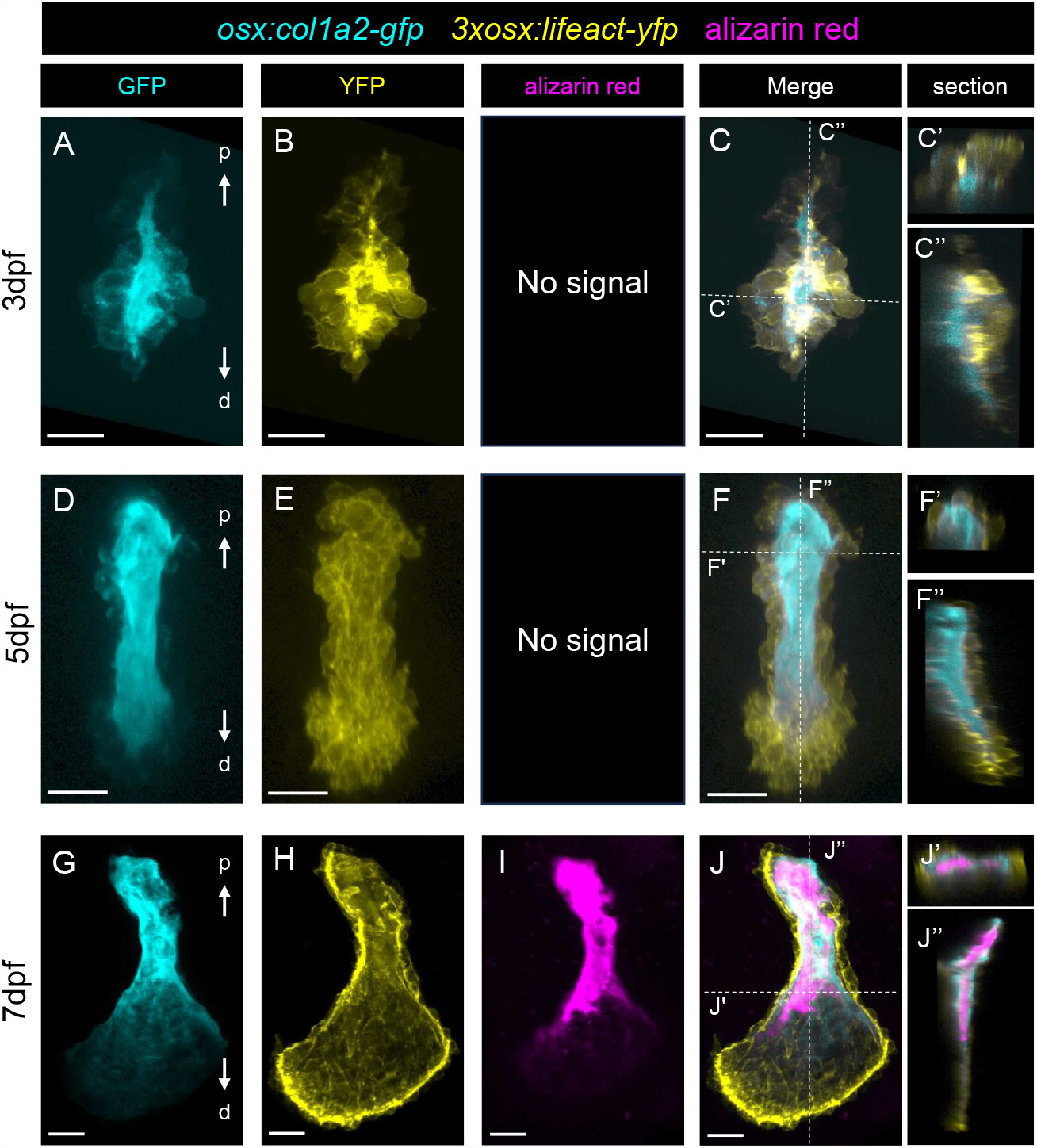
Developmental process of the opercular bone in *Tg(osx:col1a2-gfp; 3xosx:lifeact-yfp)* fish at early stage. **(A, D, G)** Collagen structures of the opercular bone at 3 dpf (A), 5 dpf (D) and 7 dpf (G) larvae. Arrow p, d indicate proximal-distal axis. **(B, E, H)** Osteoblasts forming opercular bone at 3dpf (B), 5dpf (E) and 7dpf (H). Alizarin red signal in the opercular bone. Alizarin red signal was detected in only 7dpf larvae. **(C, F, J)** Merge view of (A, B), (D, E) and (G, H, I). (C’, C’’, F’, F’’, J’, J’’) Cross-sectional views at white dotted line in (C), (F), (J). Scale bar = 20µm.

In embryos at 3 dpf, the formation of oval pouch-like osteoblasts populations was observed, and collagen structures were found in the space inside these pouch-like populations of oval osteoblasts (Fig. 5A, B, C). At 5 dpf, the collagen structures extended along proximal-distal axis while surrounded by cell populations. By 7 dpf, the opercular bone had a characteristic fan-shaped morphology (Fig. 5D, E, F, G, H, I, J). During this period, the cells surrounding collagen structure gradually shifted from an almost spherical to a flattened morphology (Fig. C’, C’’, F’’, F’’, J’’, J’’). Furthermore, strong YFP fluorescence was consistently observed throughout all developmental stages at sites where cells were in contact with collagen structures (Fig. 5C, F, J). In Tg(*3xosx:lifeact-yfp*) fish, lifeact-YFP expressed in osteoblasts visualize their actin fibers (Riedl *et al*., 2008). Thus, prominent YFP fluorescence at the cell-collagen interface indicated actin-rich regions. Furthermore, alizarin red staining revealed that calcification occurred within the bone collagen at 7 dpf (Fig. 5G, H, I, J).

## 4. Discussion

In this study, by expressing Col1a2-GFP in zebrafish under the control of an osteoblast-specific promoter, we succeeded in visualizing collagen in bones such as fin ray, opercular bones and scales specifically and clearly enough to reveal the orientation of fibers. Furthermore, we succeeded in simultaneously observing osteogenesis (calcification), collagen, and osteoblasts responsible for their synthesis in three dimensions (Fig. 1, 2, 3, and 4). This technique is a useful tool for studying the complexity of the osteogenic process and has yielded several noteworthy findings, as described below.

Both *osx* and *osc* used in this study are known osteoblast-specific marker genes (Inohaya *et al*., 2007; Renn and Winkler, 2009). The different localization of Col1a2-GFP expressed under the each promotor suggests that there may be subpopulations of osteoblast populations with different gene expression (Iwasaki *et al*., 2018), each responsible for the formation of different parts of bone which show different collagen pattern (Georgiadis *et al*., 2016; Iwasaki *et al*., 2018; Ofer *et al*., 2019). It would be important to isolate each subpopulation of osteoblasts and examine their RNA expression repertoires.

In this study, we successfully observed the dynamics of opercular bone during early development. The collagen matrix initially started to be formed in a pouch like clusters of rounded osteoblasts and then changed into a characteristic shape. At the same time, the shapes of osteoblasts surrounding it also gradually changed from spherical to flat (Fig. 5C’, C’’, F’’, F’’’, J’’, J’’), similar to observation results in previous study (Huycke *et al*., 2012). The initial round shape of the osteoblasts may be due to the absence of an extracellular matrix (collagen) to serve as a scaffold. In the previous studies, it was also reported that altered ECM structures cause morphology changes of the osteoblasts at other bones and in vitro culture assays (Kimmel *et al*., 2010; Matsugaki *et al*., 2015; Shiflett *et al*., 2019; Tsai *et al*., 2022).

Osteoblasts not only form bone collagen, but also secrete hydroxyapatite and perform bone calcification. In the observations of this study, calcification in opercular bone formation proceeded from within the collagen structure rather than from the surface, where it was in contact with the osteoblasts. This pattern of calcification was not seen in post-grown opercular bone or fin ray bone, except in regenerating fin ray bone, which calcification progressing from inside of the bone (Fig. 4A’, B’, Fig. 5I, J, J’, J’’, Sup. Fig. 2). Calcification from areas not in contact with osteoblasts suggests that osteoblasts may extend their cellular projections into the collagen and secrete hydroxyapatite from within the collagen structure (Fig. 5). More detailed observation of the dynamics of osteoblasts during bone formation may allow us to elucidate this question.

It is also important to note that the orientation of collagen can now be clearly observed in vivo along with the dynamics of osteoblasts. Since the orientation of collagen in fin ray bone is consistent with the orientation of actin fibers in osteoblasts localized in the same area (Sup. Fig. 1), it is possible that the orientation of bone collagen in fin bone is also affected by the orientation of actin fibers in osteoblasts. Indeed, microtubule and actin fiber in osteoblasts were suggested to relate collagen fiber alignment in scale in the previous study (Zylberberg *et al*., 1988). Future studies on the dynamics of osteoblast cytoskeleton and nearby collagen fibers will elucidate their relationship.

## Acknowledgements

We thank members of the Kondo laboratory for their experimental support and comments on this work. Confocal microscopy experiments were performed at the Graduate School of Frontier Biosciences Osaka University with the cooperation of Leica imaging labo and Dr. Kaneko. This work was supported by JSPS KAKENHI, Grant Number 20H05943, JSPS KAKENHI, Grant Number 21H00327, JST FOREST Program, Grant Number JPMJFR224P, Japan.

**Sup. Fig. 1 The fin ray bone in 1month old *Tg(osx:col1a2-gfp; 3xosx:lifeact-yfp)* fish**

Actin cytoskeleton in osteoblasts (YFP) and bone collagen (GFP) in fin ray bone were observed. Scale bar = 20µm.

**Sup. Fig. 2 Col1a2-GFP distribution pattern in the regenerating fin ray bone**

**(A)** The regenerating fin ray bone in *Tg(osx:col1a2-gfp)* adult fish at 12 days post amputation. **(B)** Cross-sectional images near the edge of bone segment at white dotted lines in (A). Scale bar = 20µm.

