## Supplementary figures and images for "“In Vivo Imaging of Bone Collagen Dynamics in zebrafish”"

### Sup. Fig. 1

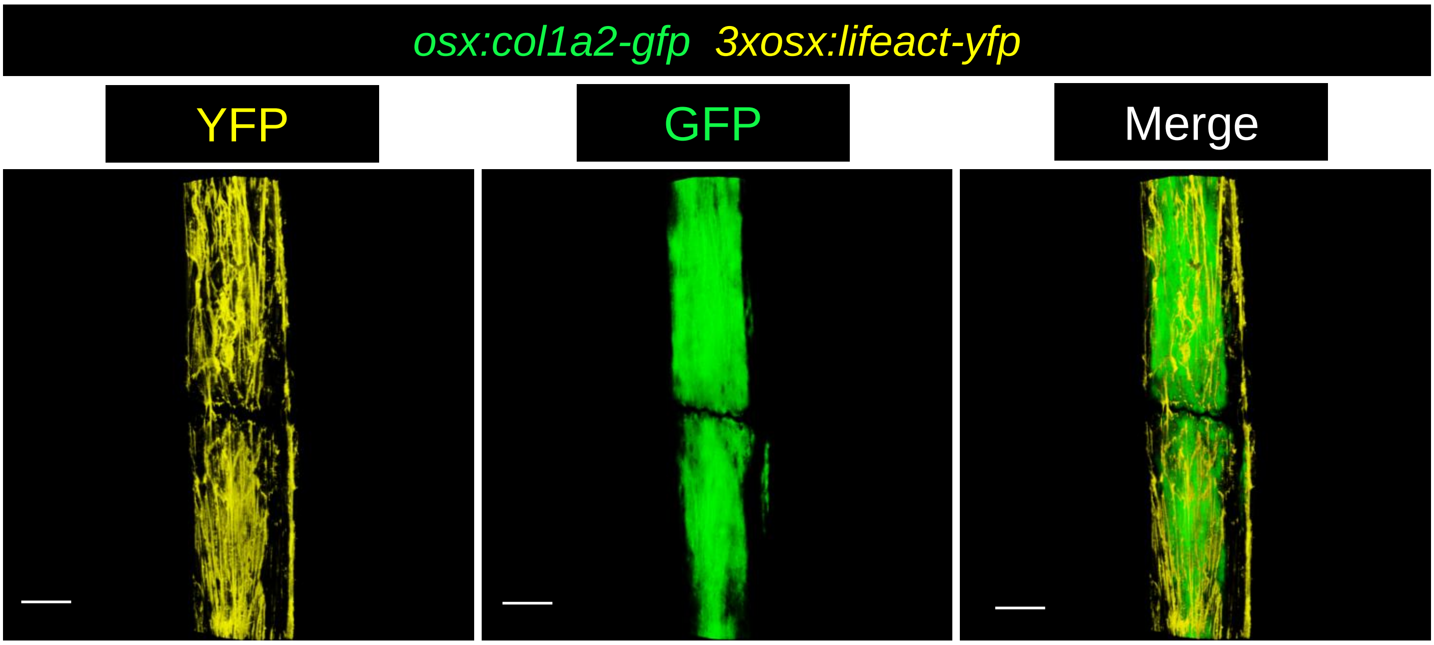

### Sup. Fig. 2

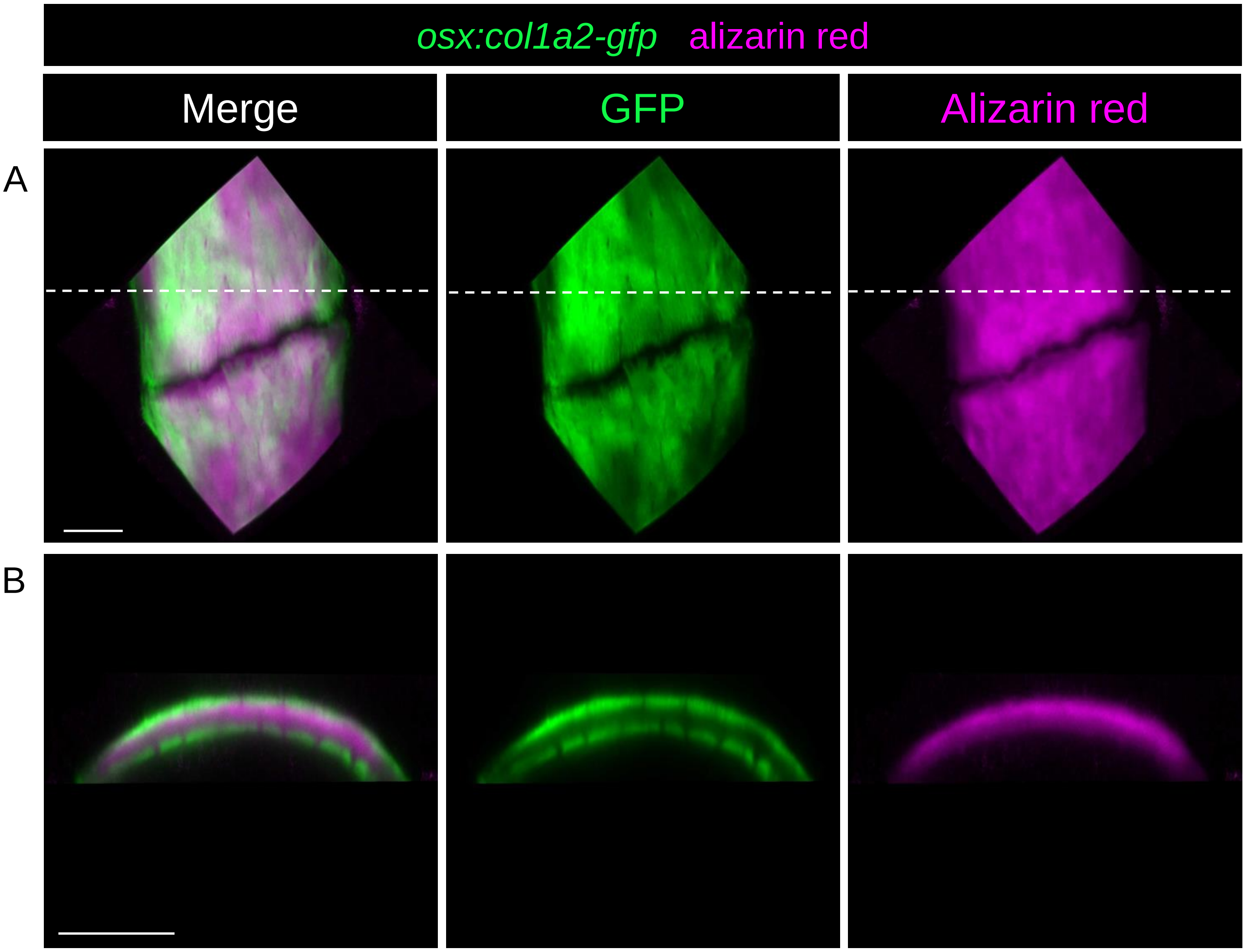
